# Efficient recovery of complete gut phage genomes by combined short- and long-sequencing

**DOI:** 10.1101/2022.07.03.498593

**Authors:** Wei-Hua Chen, Jingchao Chen, Chuqing Sun, Yanqi Dong, Menglu Jin, Senying Lai, Longhao Jia, Xueyang Zhao, Na L Gao, Zhi Liu, Peer Bork, Xing-Ming Zhao

## Abstract

Current metagenome-assembled human phage catalogs contained mostly fragmented genomes. Here, we developed a vigorous phage detection method involving phage enrichment and long-read sequencing and applied to 135 fecal samples. With ~10 times more efficient in obtaining complete genomes (~34%) than the Gut Virome Database, we identified the first megabasephage (~1.03Mb), and revealed the hidden diversity of the gut phageome including dozens of phages more prevalent than the crAssphages and Gubaphages.

## Main

The gut viral community (also known as the gut phageome), mainly consisting of bacteriophages and archaeal viruses (phages hereafter), has been shown to be diverse in the human gut^1,2^. Phages play crucial roles in shaping the gut microbial composition and hold great promise for the precision manipulation of the gut bacteriome. Despite tremendous success in identifying human (gut) phages from metagenome-assembled genomes^3–8^, the resulting phage catalogs contained mostly fragmented genomes. For example, the Gut Virome Database (GVD) that were assembled from short-read sequencing of 2,697 viral-like particle (VLP) -enriched samples contained only ~4% complete genomes^3^. Bulk-metagenomic sequencing assembled phage catalogs such as the Gut Phage Database (GPD) contained slightly higher complete genome rates (~12%), but their methods could only recover few phage genomes per sample and underestimated the diversity of the human gut phageome.

Here, we developed a vigorous phage detection method involving phage enrichment and long-read sequencing, and applied to fecal samples of 180 healthy Chinese participants. Briefly, we first used a modified VLP enrichment protocol to an increased amount of feces (~500g) to extract high-quality, high-molecular-weight (HMW) doubled stranded phage DNAs (Methods). We subjected all qualified samples to viral next-generation sequencing (vNGS) and those with sufficient amounts of HMW DNAs to PacBio third-generation sequencing (vTGS) (Fig. 1A) using the circular consensus sequencing (CCS) mode. After removing human host and bacterial contaminations, we assembled the resulting clean reads using a combined assembly strategy including vNGS, vTGS and hybrid assemblies (Methods), de-replicated at an average nucleotide identity (ANI) of 95% and obtained a total of non-redundant 97,660 contigs that were either ⩾5 kb or ⩾ 1.5 kb and circular. We filtered the contigs using six popular viral recognition tools including VirSorter^9^, VirFinder^10^and PPR-Meta^11^, and evaluated the completeness of the contigs using CheckV. We retained contigs that were either recognized as viral by two and more tools (20,444), or by one tool and of high-quality by CheckV (3,069), resulting in a catalog of 23,513 phage genomes that we referred to as the Chinese Human Gut Virome (CHGV) collection (Figure 1A).

**Figure 1.**
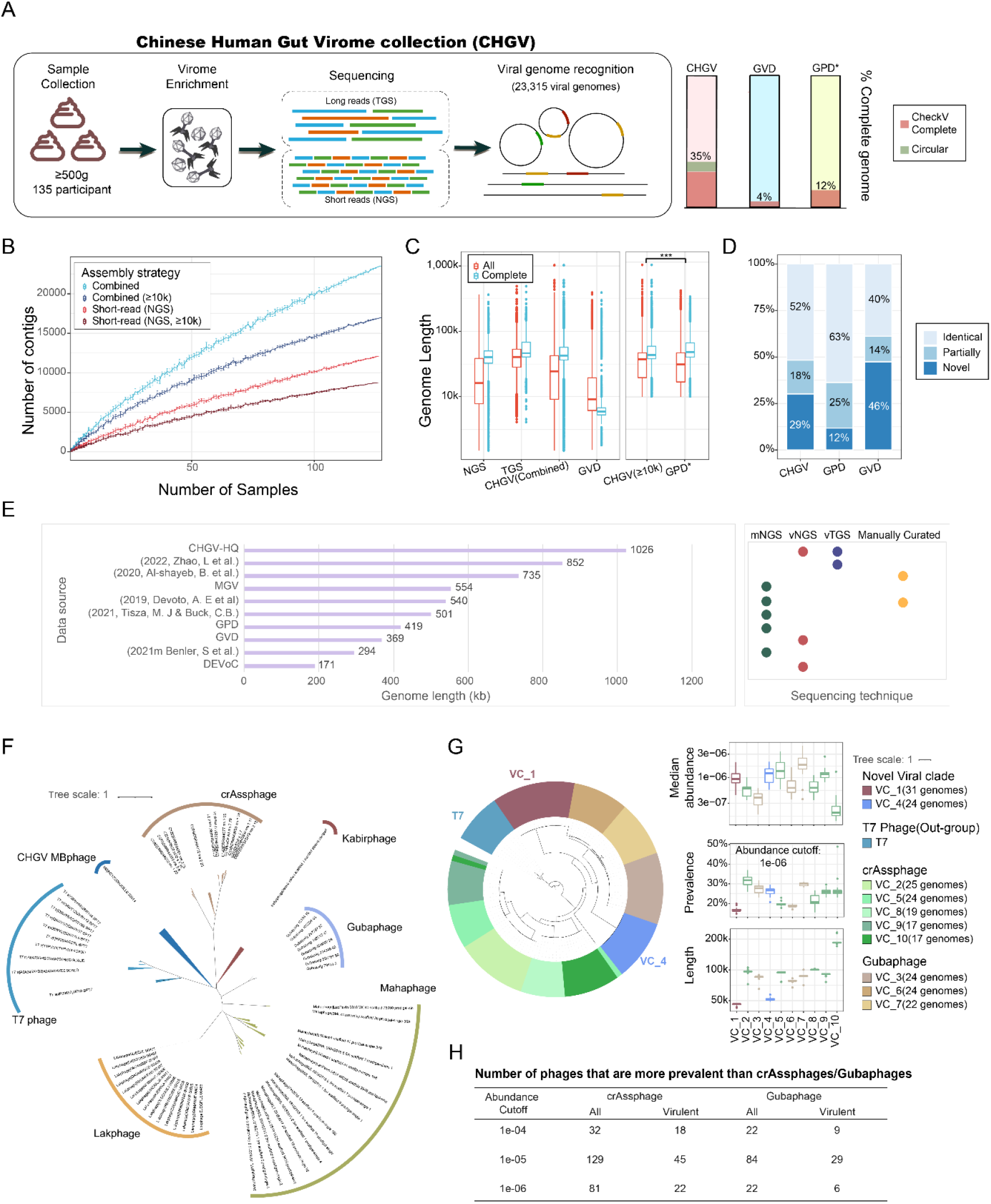
A rigorous phage detection method recovered more and longer gut phages with higher proportion of complete phage genomes. **A**, Combined assembly of long- and short reads generated a Chinese Human Gut Virome collection containing ~33% complete phage genomes. Bar plot comparing the CheckV^13^ complete(CheckV completeness 100%) genome ratio among databases. GVD: The Gut Virome Database; GPD: the Gut Phage Database. *note phage genomes <10k were excluded from the GPD catalogue. **B**, Rarefaction curves of non-redundant/unique phage contigs obtained from the short-(vNGS) and combined-assemblies, and the public VLP samples used in the GVD (vPub). **C**, Genome lengths of different assemblies and catalogues. **D**, Bar plot showing the novelty of the CHGV and selected public human viral catalogues as compared with all other human viral catalogues including GVD, GPD, CHVD^6^, DEVoC^7^, and MGV^5^. Identical: ⩾ 95% average nucleotide identity (ANI); partially: ⩾ 70% ANI; novel <70% ANI. **E**, Large phage genomes of ≥ 100 kb in size reported in recent studies^4,5,7,12,14–17^ and the corresponding identification methods. **F**, Phylogenetic relationships among the gut megaphage-1 (Gut-MBP1) identified in this study and representative phages with length ≥100 kb from public databases, including the crAssphages and Gubaphages (~100 kb), Mehaphage (~250 kb), Lakphage (~550 kb) and Kabirphage (~260 kb); T7 phages were included as the outgroups. The protein sequences of the terminase genes were used to build the phylogenetic tree using FastTree v2.1.10 ^18^. The tree was visualized using iTol^19^. **G**, Phylogenetic analysis of the top ten VCs (ranked by VC size) using terminase protein sequences (left) and their abundance and prevalence in our samples (right). An arbitrary relative abundance cutoff of 1e-6 was used to calculate the prevalence of the member phages of the VCs. **H**, Number of phages that are more prevalent than the crAssphages and Gubaphages in CHGV under different abundance cutoffs.

34.58% (8,132) of the CHGV genomes were considered complete according to either CheckV (6,348 phages) or if they were circular (3,620, Methods; see also ref.^12^), representing a 7~10 times increase in terms of the complete genome rate comparing to GVD^3^ (4%) and a 2~3 times increase comparing to GPD^4^ (12%). Our method (i.e., the combined assembly) generated more non-redundant phage genomes per sample (thus were more diverse) than NGS assembly alone (Figure 1B) and longer genomes than GVD (and GPD under the same length filtering criteria, i.e. >10k; Figure 1C). In addition, our CHGV catalog included 29% novel phage genomes (i.e., those that shared <75% ANI with public viruses; Method) that were not found in any published human virome datasets including the GVD, GPD, Cenote-Taker 2–compiled Human Virome Database (CHVD)^6^, Danish Enteric Virome Catalog (DEVoC)^7^, and Metagenomic Gut Virus catalog (MGV)^5^, which is significantly higher than GPD (~12% novel genomes). GVD contained higher proportion of novel phages (46%, Figure 1D), likely because of its significantly large sample size (2,697 VLP-samples).

Our method identified the first megabase phage (MBphage), Gut-MBP1 of 1,026 kb in size that was larger than any bacteriophages ever reported^4,5,7,12,14–17^ (Fig. 1E). Phylogenetic analysis using the protein sequences of the terminase genes indicated that the Gut-MBP1 formed its own clade from other known large phages (Fig. 1F). We identified additional 20 potential MBphages in CHGV and GVD using the terminase proteins (Supplementary, Methods). These phages ranged from 5 kb~ 536 kb in size, likely being only fragments; among them, three showed high overall protein similarities with Gut-MBP1. Our data thus extended the upper limit of phage genome size and blurred the boundary between the living and nonliving.

Our method also revealed the hidden diversity of the human gut phageome in the following two aspects. First, by grouping the CHGV genomes into 1,982 non-singleton viral clusters (VCs) using the Markov clustering algorithm^20^ (Methods), we identified a VC_1 that was more diverse (i.e., contained more phage genomes) than all the VCs corresponding to crAssphages and Gubaphages, the two known most diverse phage clades in the human gut^4^, and a VC_4 that was more diverse than most other VCs (Figure 1G). Both VCs contained novel phages that were not found in the NCBI Viral RefSeq database, and formed their own clades in a phylogenetic tree consisting the genomes in the top 10 VCs. Their members were highly abundant (comparable to that of the crAssphages and Gubaphages) and prevalent (with a median prevalence of 16.3% and 26.6% at an arbitrary relative abundance cutoff of 1e-6, respectively) in our samples, and were of 40 and 50 kb in size (Fig. 1G). Additional 81 potential VC_1 phages could be identified in the GPD and MGV databases^3–7^ that shared high overall sequence similarities (Supplementary). Second, we identified at least dozens of phages that were more prevalent than the most prevalent crAssphages and Gubaphages in our samples, regardless of the relative abundance cutoffs (Fig. 1H).

In summary, our rigorous phage detection method was highly efficient in recovering complete phage genomes from human feces and significantly expanded our knowledge on the hidden diversity of the human gut phageome in multiple dimensions.

## Methods

### Sample collection

Human fecal samples were obtained from healthy volunteers recruited in Wuhan and Shanghai, China. All volunteers remained anonymous but were asked to complete a questionnaire to collect relevant information such as their sex, age, height, weight, health status, and recent antibiotic usage (Table S1). The exclusion criteria included (1) the use of antibiotics or probiotic supplements up to one month before the study; (2) the use of drugs known to significantly affect the gut microbiota composition, such as metformin^1,2^, statin^3^ or proton-pump inhibitors^4,5^, in the month prior to sample collection; (3) current chronic intestinal diseases or a history of intestinal diseases; and (4) menstruation at the time of sampling in females. After collection, the samples were immediately cooled with dry ice and transferred to a −80°C freezer within five hours. To obtain a large amount of feces for phage extraction, up to three stool samples were collected from each participant and mixed together; the mixed samples totaling at least 500 grams were processed further. In total, 163 qualified samples were obtained (Table S1).

This study was approved by the Ethics Committee of the Tongji Medical College of Huazhong University of Science and Technology, Wuhan China (No, S1241) and the Human Ethics Committee of the School of Life Sciences of Fudan University, Shanghai China (No, BE1940).

### Virome enrichment and short- and long-read sequencing

The virome enrichment protocol applied to the fecal samples was adapted from ref.^6^ with modifications to accommodate the large quantity of the collected feces from each participant. Briefly, 400~500 g of frozen feces taken from a −80°C freezer was added to five liters of SM (200 mM NaCl, 10 mM MgSO4, 50 mM Tris-HCl (pH 7.5)) buffer and stirred by an automated stirrer (A200plus, OuHor, Shanghai, China) at low speed (120 rpm) at room temperature until all feces were dispersed. Then, the suspended mixture was filtered through four layers of gauze (21 s x 32 s/28 × 28) and centrifuged at 5000 x g for 45 min at 4 °C. The supernatant was transferred to fresh tubes and centrifuged at 8000 x g for 45 min at 4 °C. The supernatant was subsequently concentrated to ~300 ml via a 100 KD ultrafiltration membrane (Sartorius, VIVO FLOW 200). NaCl was then added to the filtrates to a final concentration of 0.5 mol/L, and the samples were stored at 4 °C for one hour. Then, PEG 8000 was added to a final concentration of 10% w/v, and the samples were incubated at 4 °C overnight. On the following day, phage particles were sedimented at 13000 x g for 35 min at 4 °C.

The obtained pellets were fully suspended in 18~36 mL TE buffer and treated by gently shaking with an equal volume of chloroform. The mixture was centrifuged at 3500 x g for 10 min at 4 °C. The aqueous phase was then transferred to a sterile round-bottomed flask and evaporated for 15 min using a rotary evaporator at room temperature to remove traces of chloroform, which could affect the activity of DNase I in the subsequent step. The aqueous phase was transferred to a new centrifuge tube, TE buffer was added to recover the volume before treatment with chloroform, and DNase buffer was added to a 1× final concentration. Then, for every 6 mL of supernatant, 50 μL of a DNase I mixture (33.3 U/μL, Biolab) and 25 μL of an RNase A mixture (0.5 U/μL, Biolab) were added, and the resultant mixture was incubated in a thermostatic oscillator (THZ-C, Peiying, Suzhou, China) at 100 rpm for 30 min at 37 °C before the enzymes were inactivated by the addition of EDTA buffer (final concentration 35 mM) and incubation at 70°C for 10 min.

Nucleic acid was then extracted using a HiPure HP DNA Maxi Kit (D6322, Magen, Guangzhou, China) according to the manufacturer’s instructions. Briefly, proteinase K and SDS lysis buffer were added, and the mixture was then incubated at 56 °C for one hour. Viral particles were further lysed by adding the CFL buffer provided with the kit, and the lysates were subsequently treated with an equal volume of phenol:chloroform:isoamyl alcohol (25:24:1, pH 8.0), followed by centrifugation at 12000 × g for 15 min at room temperature. After centrifugation, the supernatant was transferred to a new centrifuge tube and treated with an equal volume of chloroform with gentle shaking, followed by centrifugation at 12000 x g for 15 min at room temperature. The aqueous phase was transferred to a new tube, loaded onto a DNA Mini Column provided by the kit, and centrifuged at 12000 x g for 1 min. The DNA Mini Column was then washed with GDP and GW2 buffers. DNA was eluted using DNA elution buffer and stored at −80 °C for further analysis. Note that all buffers and columns used in this part of the study were provided in the kit.

The purified VLP DNAs were quality checked and subsequently sequenced on the Illumina (short-read) and PacBio (long-read) platforms. For Illumina sequencing, nucleic acids were sheared with a g-TUBE (Covaris, USA) to generate a target size fragment of 400 bp, followed by sequencing library construction using the Nextera XT DNA Library Preparation Kit (Cat. No. FC-131-1096, Illumina, USA) according to the manufacturer’s instructions and sequencing using an Illumina HiSeq2000 sequencer (Novogen, Beijing, China) to generate paired-end reads of 150 bp. The generated dataset was then referred to as viral next generation sequencing (vNGS) data. For PacBio sequencing, DNAs were sheared into approximately 5 kb fragments by using a g-TUBE (Covaris, USA) and purified with AMPure PB magnetic beads, followed by a quality check using 0.7% agarose gel electrophoresis. The qualified samples were employed to construct sequencing libraries using the SMRTbellTM Express Template Prep Kit 2.0 (Pacific Biosciences, USA) according to the manufacturer’s instructions. The quality of the DNA libraries was checked with an Agilent 2100 Bioanalyzer (Agilent Technologies, USA), and the libraries were then sequenced with a PacBio RS II sequencer (Pacific Biosciences, Menlo Park, CA, USA) in circular consensus sequencing (CCS) mode. The generated dataset was then referred to as viral third generation sequencing (vTGS) data.

### Raw data processing

Raw next generation sequencing of viral reads (referred to as vNGS hereafter) were processed with Trimmomatic v0.38^7^ (with parameter LEADING:3 TRAILING:3 SLIDINGWINDOW:15:30 MINLEN:50) to remove adaptors and trim low-quality bases; reads of 50 bp or less after trimming were discarded. The third generation sequencing of viral reads (referred to as vTGS) reads were corrected with CCS using pbccs (v4.0.0, https://github.com/nlhepler/pbccs) with the default parameters.

Putative human reads were identified from the trimmed/CCSed reads by aligning the latter to the human reference genome (hg38; GCA_000001405.15) using Bowtie2^8^ (v2.4.2, --end-to-end) with default parameters and removed from further analysis.

In total, we obtained 4.89 terabytes of clean data for the vNGS samples and 561 gigabytes of CCSed data for the vTGS samples.

### Combined assembly of short- and long-reads

Briefly, IDBA-UD^9^ (Release 1.1.3, parameters: --maxk 120 --step 10 –min_contig 1000) was used to assemble the filtered vNGS data. Canu^10^ (v2.0-, parameters: genomeSize=20k corOutCoverage=1 -corrected) and Flye^11^ (v2.8.2, parameters: --meta --genome-size 20k --min-overlap 1000) were used to assemble the filtered vTGS CCS reads. Because Canu does not have a meta-assembly mode and tends to extend contigs by merging DNA sequences from different viral species to generate erroneous contigs, unitigs were used for subsequent analysis; unitigs are basic blocks of contigs that are shorter but more reliable than contigs (‘unitigs’ are derived from contigs; wherever a contig end intersects the middle of another contig, the contig is split)^12^. To further extend the sequences, MetaBAT2^13^ (version 2, default parameters) was used to group unitigs into bins. If all unitigs from one contig could be grouped into the same bin, contigs instead of unitigs were used for further analysis. OPERA-MS^14^ (v0.9.0, parameters: -contig-len-thr 1000 --polishing --no-strain-clustering --no-ref-clustering) and metaSpades^15^ (v3.13.1, default parameters) were employed for hybrid assemblies using both the vTGS and vNGS datasets from the same samples(Figure S1).

Contigs/unitigs obtained from all the above three strategies were merged; for samples that did not have vTGS data, contigs from the IDBA-UD assembler were used.

The merged dataset was dereplicated using CD-HIT^16^ (v4.8.1, parameters: -c 0.95 -n 8) using a global identity threshold of 95%.

### Prediction of viral contigs with state-of-the-art tools

To identify viral contigs, six independent state-of-the-art viral identification pipelines were used, including VirSorter v2.0^17^ (--min-score 0.7), VirFinder v1.1^18^ (default parameters), and PPR-Meta v1.1^19^ (default parameters). A BLAST search against the Viral RefSeq genomes was also performed using BLASTn v.2.7.1^20^ with the default parameters and an *E*-value cutoff of <1e-10; Release 201 (Jul 06, 2020) of the Viral RefSeq database contained 13,148 viral genomes. In addition, the annotated protein sequences were used for BLAST searches against the NCBI POG (Phage Orthologous Groups) database 2013^21^.

A contig was annotated as a virus if it was circular/met at least two out of the following criteria 1-5, adopted from the Gut Virome Database (GVD) ^22^:

- VirSorter score ⩾ 0.7,
- VirFinder score > 0.6,
- PPR-Meta phage score > 0.7,
- Hits to Viral RefSeq with > 50% identity & > 90% coverage,
- Minimum of three ORFs, producing BLAST hits to the NCBI POG database 2013 with an *E*-value of ⩽ 1e-5, with at least two per 10 kb of contig length.
- Alternatively, contigs met one of the above criterium and were annotated as high-quality (≥ 90% completeness) by CheckV^23^ were also annotated as viruses.

As short contigs may only represent fragments of viral genomes, contigs that were longer than 5 kb or circular contigs longer than 1.5 kb were selected for further analyses; this dataset was referred to as the Chinese Human Gut Virome (CHGV) dataset, which consisted of a total of 23,513 viral populations.

Rarefaction curves were generated by randomly resampling the pool of N samples 10 times and then plotting the number of dereplicated (unique) contigs found in each set of samples.

### Public viral genome databases/catalogs used in this study

The following public human virome databases were used in this study. GPD, the Gut Phage Database^24^, includes 142,000 viral genomes assembled from metagenome sequencing. GVD, the gut virome database^22^, includes 33,242 viral genomes assembled from Viral like particles (VLP) sequencing. MGV, the Metagenomic Gut Virus collection^25^, includes 54,118 candidate viral species assembled from metagenome sequencing. CHVD, the Cenote Human Virome Database^26^, includes 45,033 viral taxa assembled from metagenome sequencing. DEVoC, the Danish Enteric Virome Cataloge^27^, includes 12,986 viral genomes assembled from VLP sequencing. The NCBI viral Reference genomes, Release 201 (Jul 06, 2020) of the Viral RefSeq database contained 13,148 viral genomes.

### Identification of complete phage genomes in CHGV and public viral datasets

The CheckV^23^ program were used on the CHGV and public viral datasets, those that were annotated with 100% completeness were considered to be complete genomes (CheckV complete).

In addition, a customized pipeline was used to identify circular contigs that were considered as complete genomes in CHGV. First, the BLASTn program^20^ was used to search for alignable regions within each contig; if the front and tail portions of the contig were exact matches over 30 base pairs (nucleotide identity=100, *E*-value<1e-5), they were considered as circular genomes^28^. Second, the clean sequencing reads were mapped to the CHGV genomes using either pbmm2 (https://github.com/PacificBiosciences/pbmm2) for the vTGS data or bowtie2^8^ for the vNGS. Genomes with at least two reads mapped to both the front and tail of the genome were considered to be circular genomes, resulting additional 1,054 circular genomes.

### Estimating the proportion of novel viral genomes in one dataset as compared with all other viral databases

To estimate the proportion of novel viral genomes in one dataset, the BLASTn tool was used to search all its sequences against all other viral databases mentioned above. Average nucleotide identity (ANI) was calculated by merging the hit regions with identity ⩾90%, and hit length ⩾ 500bp, then calculated the coverage of these regions. Based on the overall ANI, a viral sequence is considered to be identical, partial identical or novel if it has ⩾ 95%, ⩾ 70% or <70% ANI as compared with other viral sequences.

### Identification of the first megabasephage (MBphage)

We extracted the longest genome with ~1,026 kb in length. Gut-MBP1 was classified as high-quality by CheckV^37^ (Methods). The annotation with VirSorter2 excluded the possibility of a nucleocytoplasmic large DNA virus (i.e., NCLDV; Methods).

We observed overall even sequencing-depth of 1,740× and 412× across the Gut-MBP1 genome according to the vNGS and vTGS reads, respectively, thus excludes the possibility of false assembly. A peak region of ~5.8kb was found between 745~751 kb was presumably due to repeated sequences; the assembly of this region was supported by 58 vTGS reads that covered the whole region and ~1,000 vTGS reads that spanned the junctions on both sides of the region (~470 vTGS reads at each side, Figure S2).

### Identification of longest viral genomes from public databases and literature

Longest bacterial phages were extracted from recent studies including 2022. Zhao, L *et al*.^29^; 2020, Al-Shayeb, B. *et al*.^30^; 2019, Devoto, A. E. *et al*.^31^; 2021, Tisza, M. J. & Buck, C. B.^32^; 2021, Benler, S. *et al*.^28^, and public databases including MGV ^25^, GPD ^24^and GVD ^22^. DeVoc.

### Functional annotation of CHGV proteins

The encoded protein sequences of the CHGV genomes were annotated using Prodigal ^33^ v2.6.3 with default parameters.

Proteins translated from the CDS sequences were then annotated with eggNOG mapper v1.0.3-3^34^ and hmmscan^35^ v3.3.2 against Pfam^36^ v34.0, and VOGdb v204 (*E* value <1e-5, score >=50, http://vogdb.org/).

The terminase protein sequences were extracted to conduct phylogenetic analysis (below section).

### Phylogenetic analysis of selected phages

Phylogenetic analysis was performed for selected phages using the terminase protein sequences. Briefly, for each group of phages of interest, their terminase protein sequences were aligned using MUSCLE^38^ v3.8.1551 with the default parameters. Phylogenetic trees were built with FastTree^39^ v2.1.10 with default parameters. Phylogenetic trees were then visualized and annotated using iTol^40^ and EvolView^41^.

Representative phages with length ⩾100 kb were obtained from the following public datasets, including the crAssphages^42^ and Gubaphages^24^ (~100 kb), Mahaphage^30^ (~250 kb), Lakphage^31^ (~550 kb) and Kabirphage^30^ (~260 kb).

### Clustering viral contigs into viral clusters (VCs)

The clustering of gut viral contigs into viral clusters (VCs) was performed using a strategy adopted from the GPD^24^. Briefly, a BLASTn algorithm with default parameters was used to search the nucleotide sequences of the CHGV viral contigs against themselves for homologous sequences. An *E*-value threshold of 1E-10 was first used to filter the BLASTn results; the BLASTn query-hit pairs were further filtered to retain those with a coverage > 70% on larger genomes and coverage >90% on smaller genomes. Here, the coverage was calculated by merging the aligned fraction length of BLASTn high-scoring pair (HSP) sequences that shared at least 90% nucleotide similarity. Finally, a Markov clustering algorithm^43^ (MCL v14-137) was used with an inflation value of 4.0, which took the filtered BLASTn results as input, carried out graph-based clustering and clustered the viral contigs into VCs.

### Identification of crAssphages and Gubaphages in CHGV contigs

crAss-like phage genomes were annotated by following the method reported in a previous study ^44^. First, the nucleotide sequences of all CHGV contigs were subjected to search against the protein sequences of the polymerase (UGP_018) and the terminase (UGP_092) of the prototypical crAssphage (p-crAssphage, NC_024711.1) using BLASTx. Second, the nucleotide sequence similarities between the CHGV contigs and the p-crAssphage genome were assessed using BLASTn. A contig was then labeled as a putative crAssphage when it was longer than 70 kb and met at least one of the following criteria:

1. BLASTx hit with an *E*-value <1e-10 against either p-crAssphage polymerase or terminase
2. ⩾95% nucleotide identity over 80% of the contig length with the p-crAssphage genome

Gubaphage genomes were annotated by clustering viral contigs with the Gubaphage genomes obtained from the GPD database^24^ into viral clusters (VCs) using the methods mentioned above. Viral contigs that were in the same VC as Gubaphage were annotated as Gubaphages.

### Estimation of the relative abundance of the CHGV genomes at the viral contig and VC levels

To estimate the abundance of viral contigs, the vNGS clean reads were mapped to the CHGV database using Bowtie2. Then, we calculated the reads per kilobase million (RPKM) value of each viral contig. Relative species abundance was calculated by dividing the RPKM of a specific viral contig by the total RPKM of all viral contigs.

To avoid the noise of low-abundance taxa, viral contigs with relative abundances lower than 0.0001% were removed and the relative abundances were recalculated so that the total abundances of all contigs were added to 100%.

VC abundance was generated based on viral contig abundances by dividing the sum of the RPKM values of the viral contigs from the same viral cluster by the total RPKM value.

## Funding

This research is supported by National Natural Science Foundation of China (32070660 to W.H. C; 61932008, 61772368 to X.M. Z; 31770132, 81873969 to Z. L), National Key Research and Development Program of China (2020YFA0712403 to X.M. Z; 2019YFA0905600 to W.H. and Z.L.) and Shanghai Municipal Science and Technology Major Project (2018SHZDZX01 to X.M. Z).

## Ethics approval

This study was approved by the Ethics Committee of Tongji Medical College of Huazhong University of Science and Technology (No, S1241) and the Human Ethics Committee of the School of Life Sciences of Fudan University (No, BE1940).

## Author contributions

WHC, XMZ, ZL and PB designed and directed the research; JC managed the sampling and performed most of the experiments; CS performed most of the analysis; XZ and MJ also helped with the sample collection and phage enrichment experiments; YD performed the machine learning analysis. CS and JC wrote the paper with results from all authors; WHC, XMZ, ZL and PB polished the manuscript through multiple iterations of discussions with all authors. All authors have read and approved the final manuscript.

## Notes

### Competing Interest Statement

The authors have declared no competing interest.

### Summary of Updates

Results, figures revised. Supplemental files updated.

