## Supplementary figures and images for "Efficient recovery of complete gut phage genomes by combined short- and long-sequencing" (from another version of this preprint: version 1, 2022-07-04)

### Supplementary Figure 1

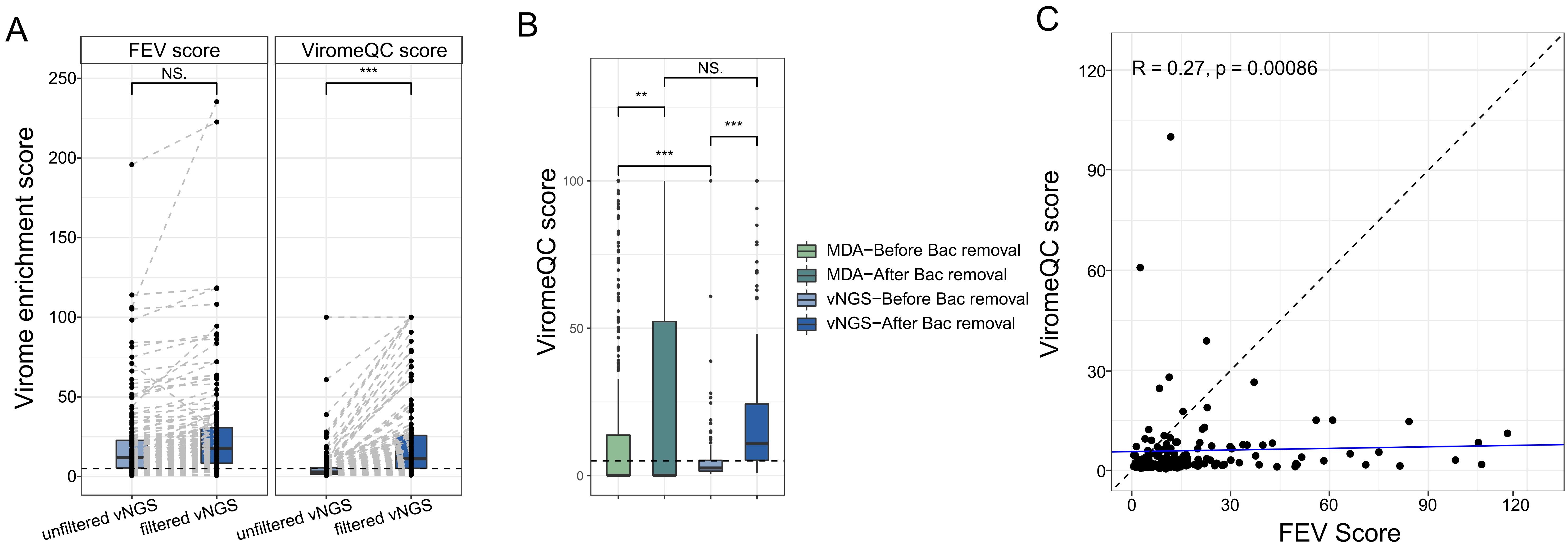

### Supplementary Figure 2

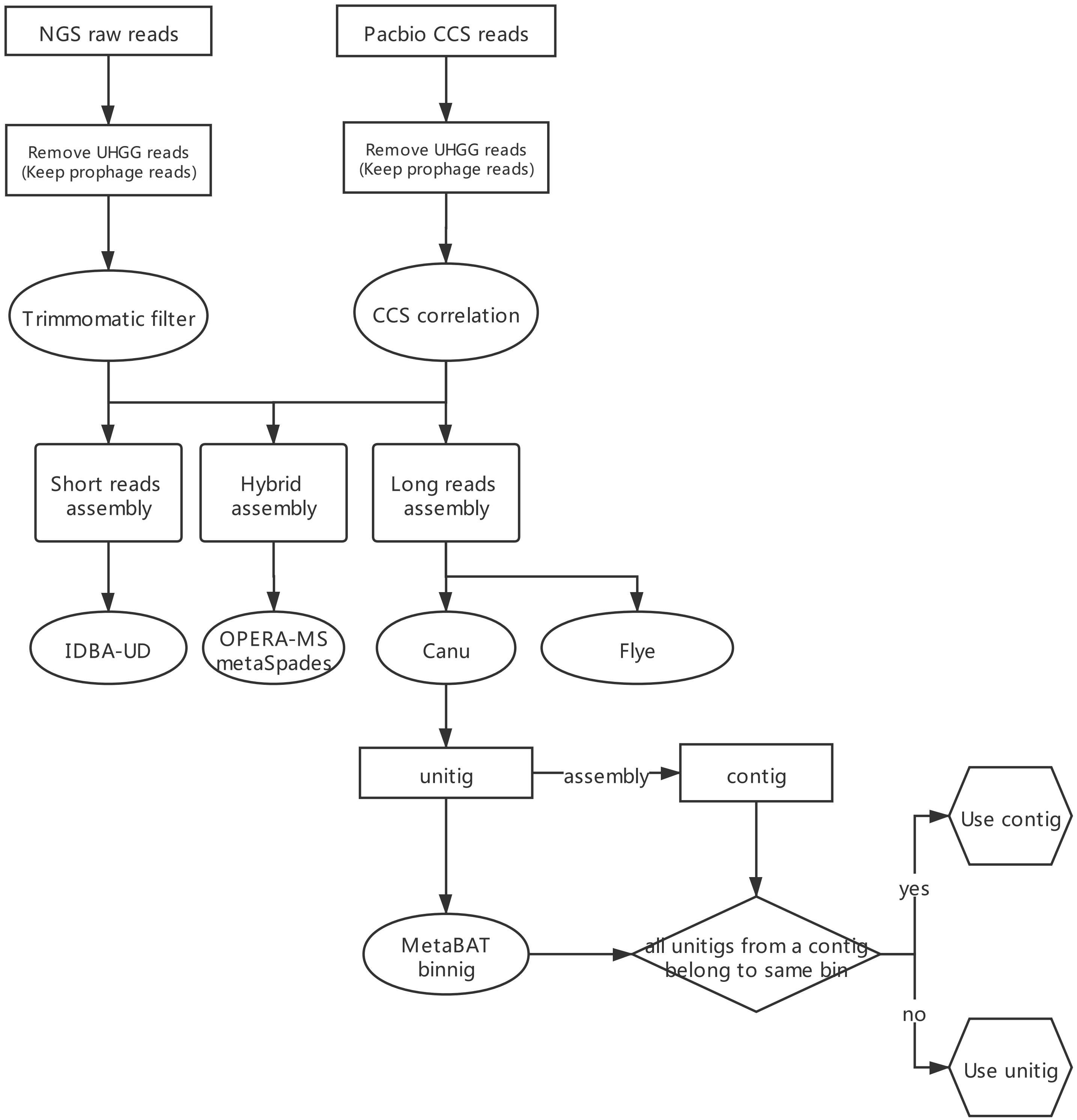

### Supplementary Figure 3

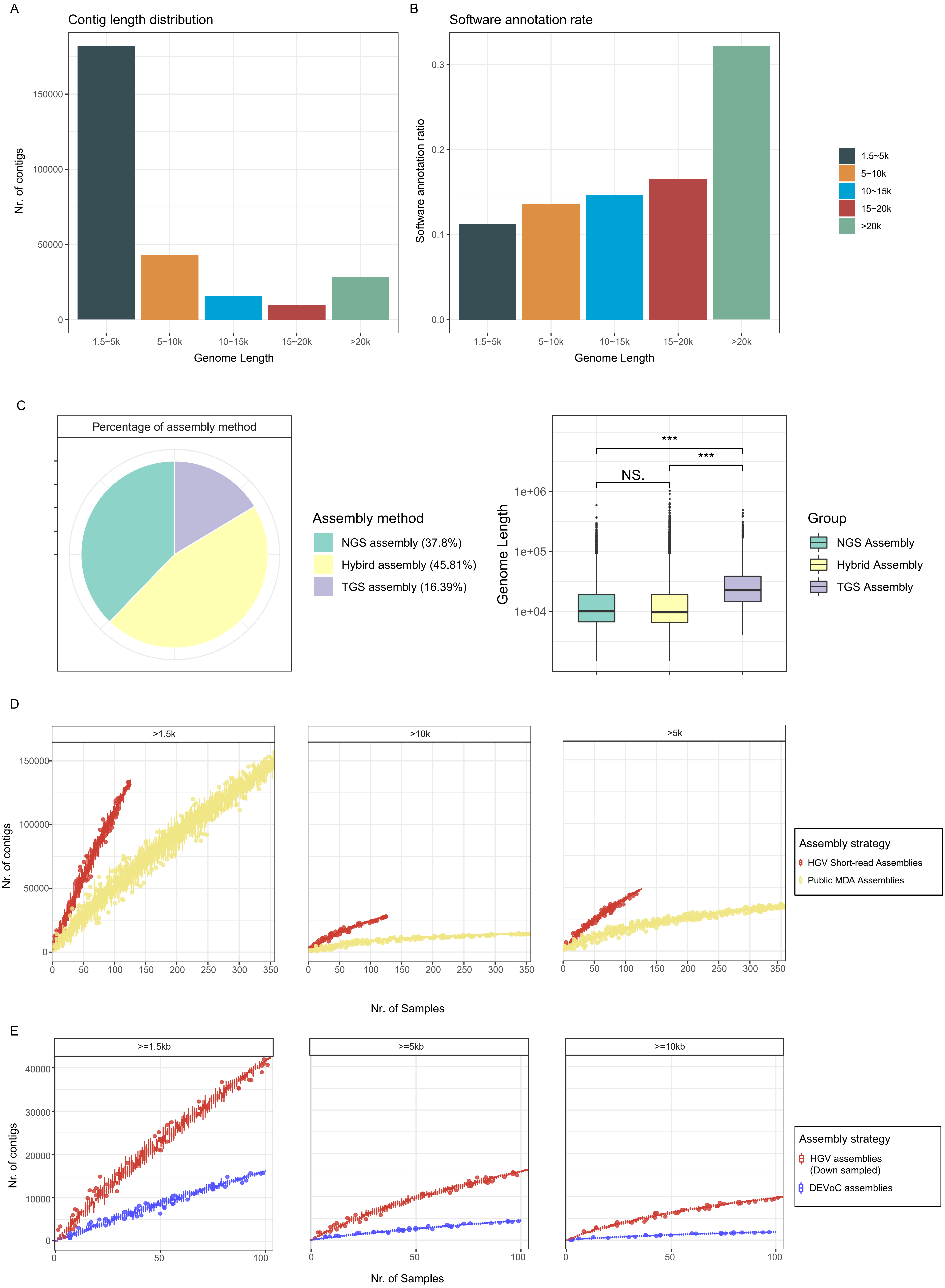

### Supplementary Figure 4

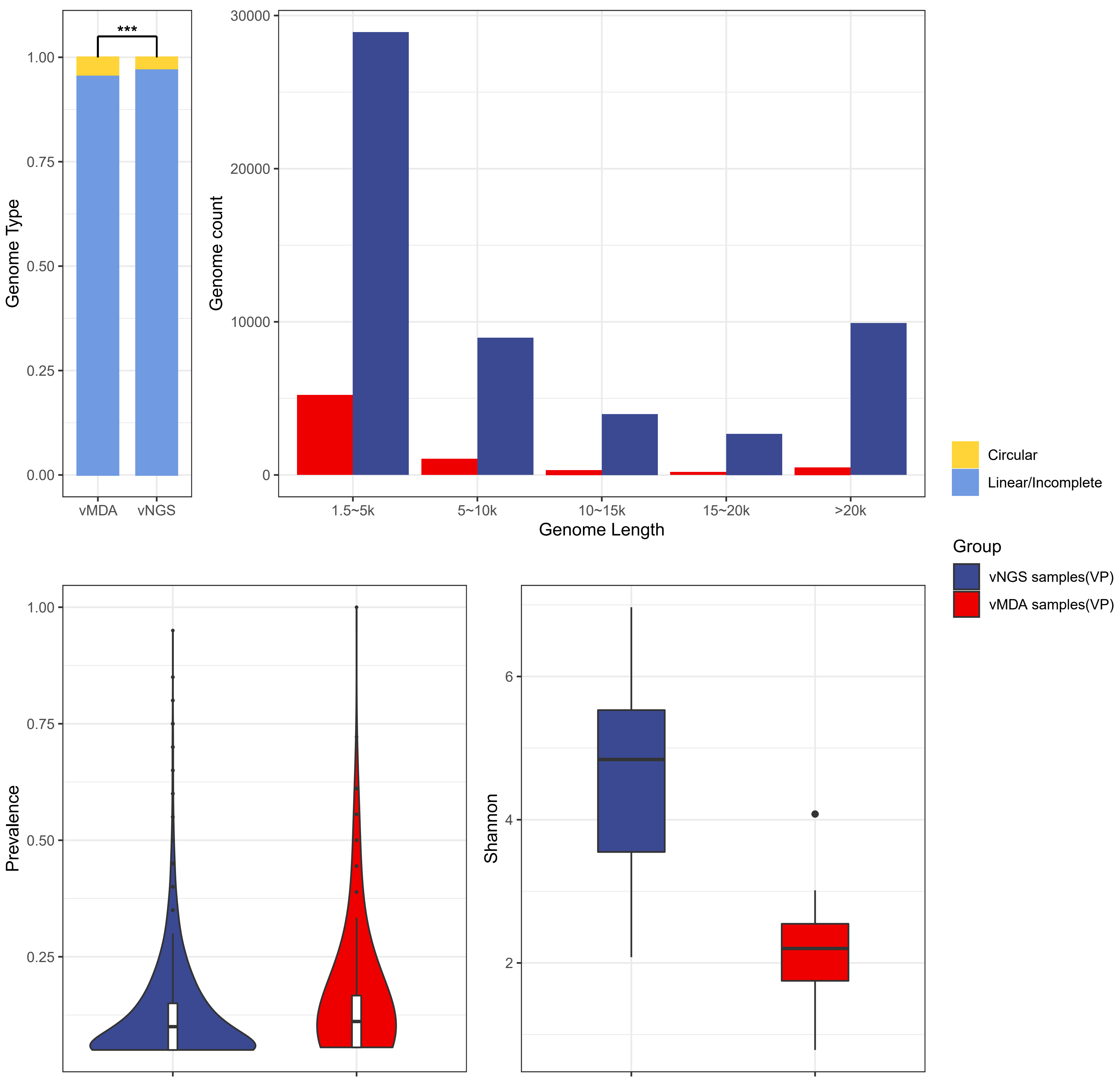

### Supplementary Figure 5

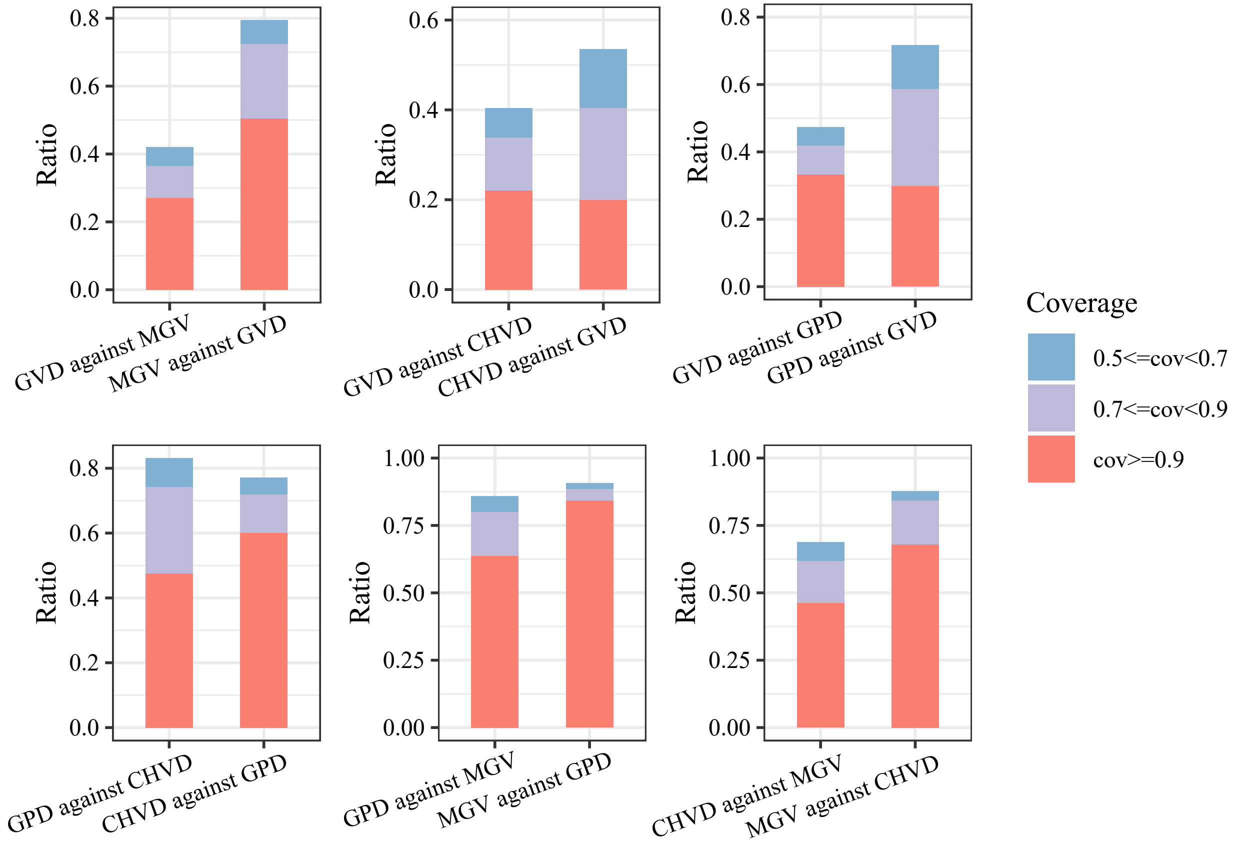

### Supplementary Figure 6

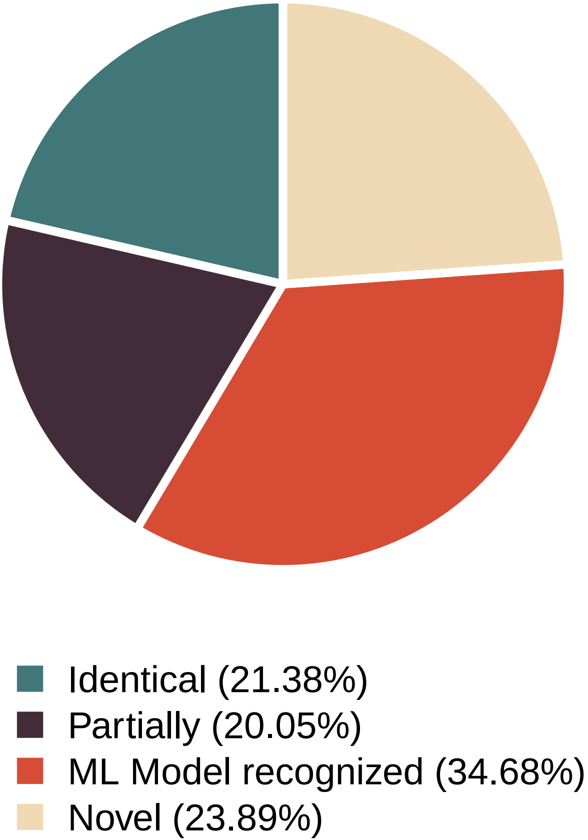

### Supplementary Figure 7

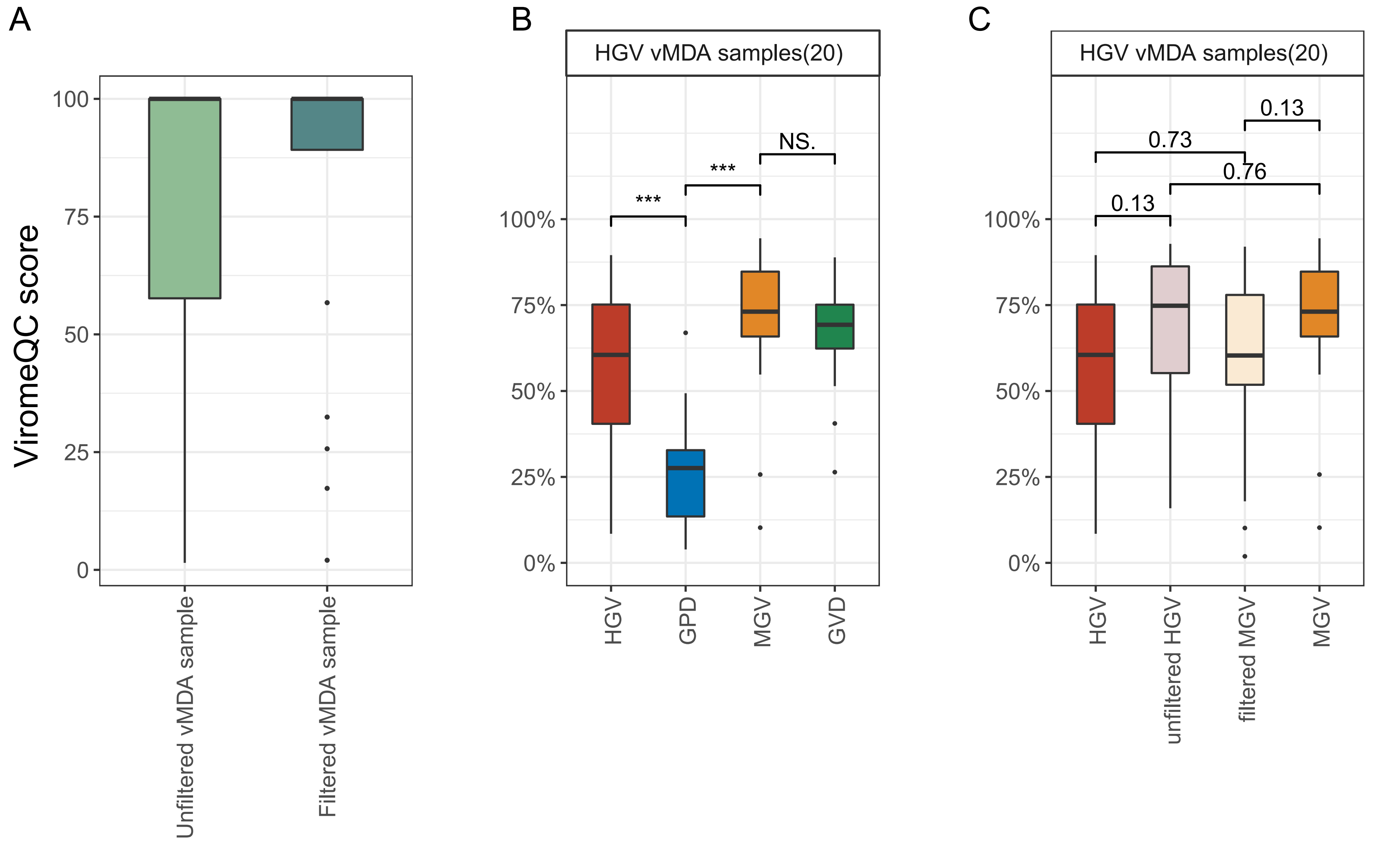

### Supplementary Figure 8

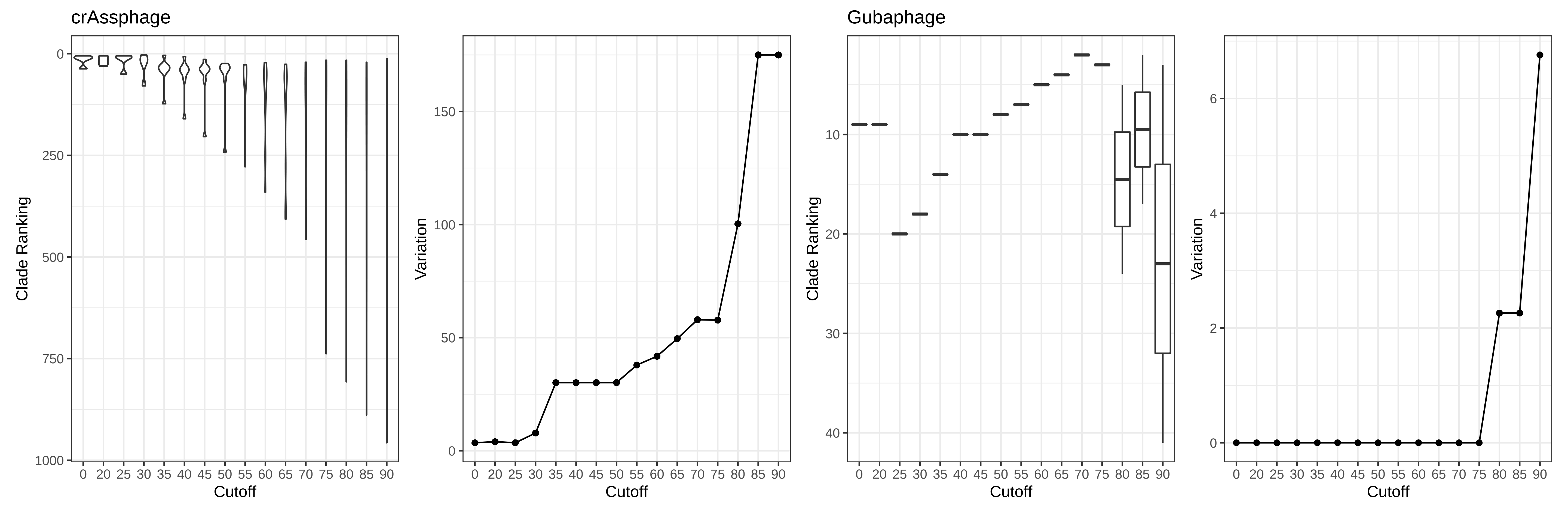

### Supplementary Figure 9

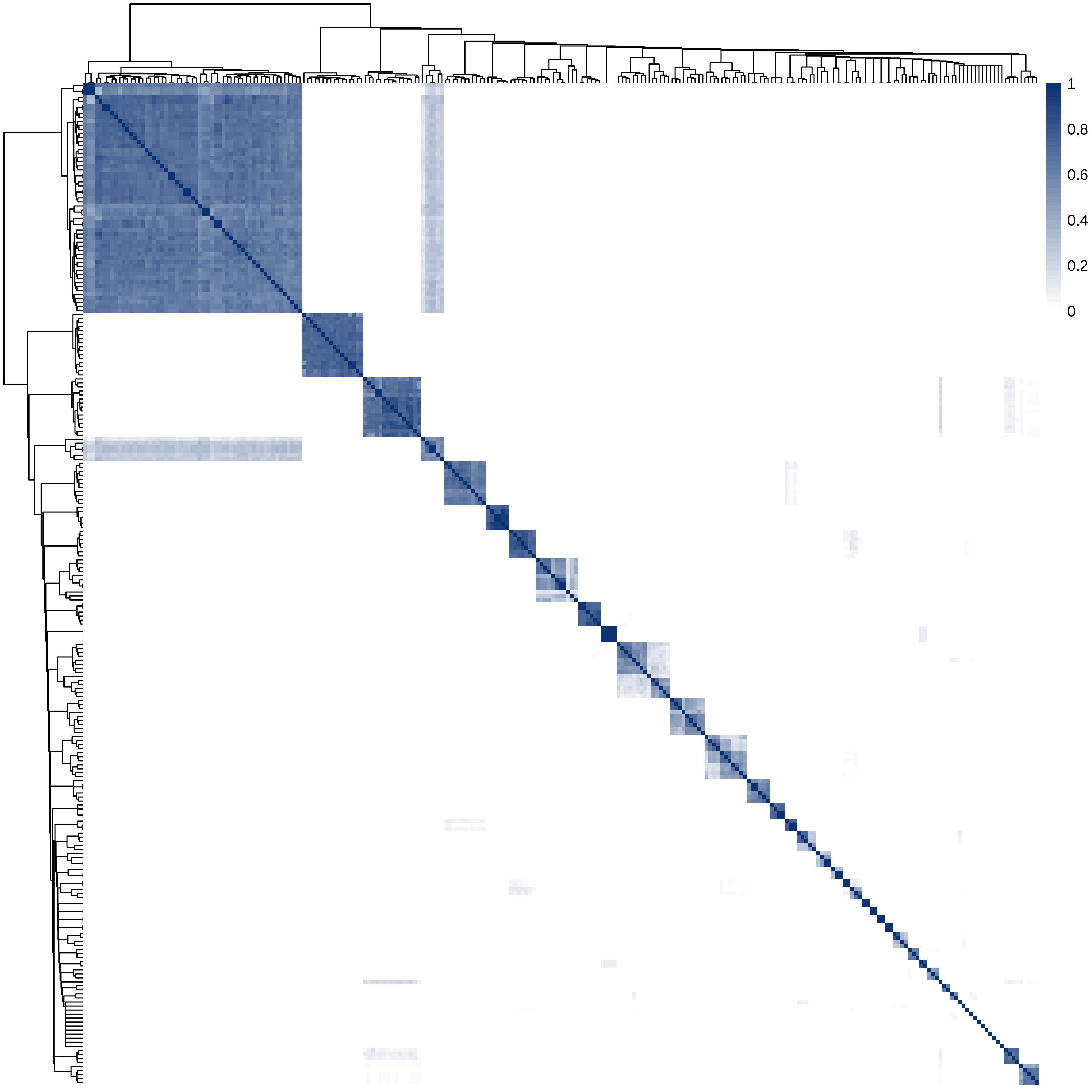

### Supplementary Figure 10

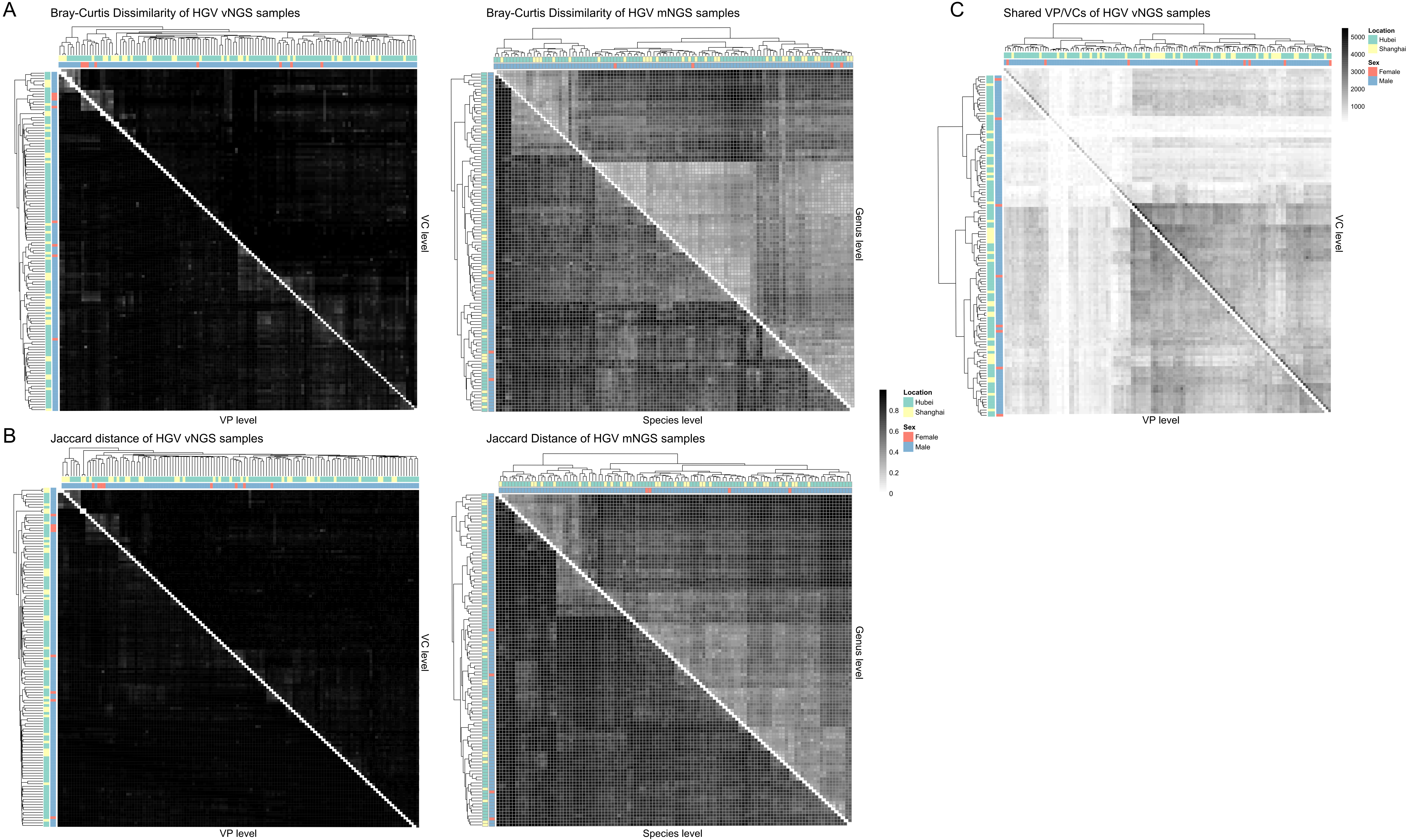

### Supplementary Figure 11

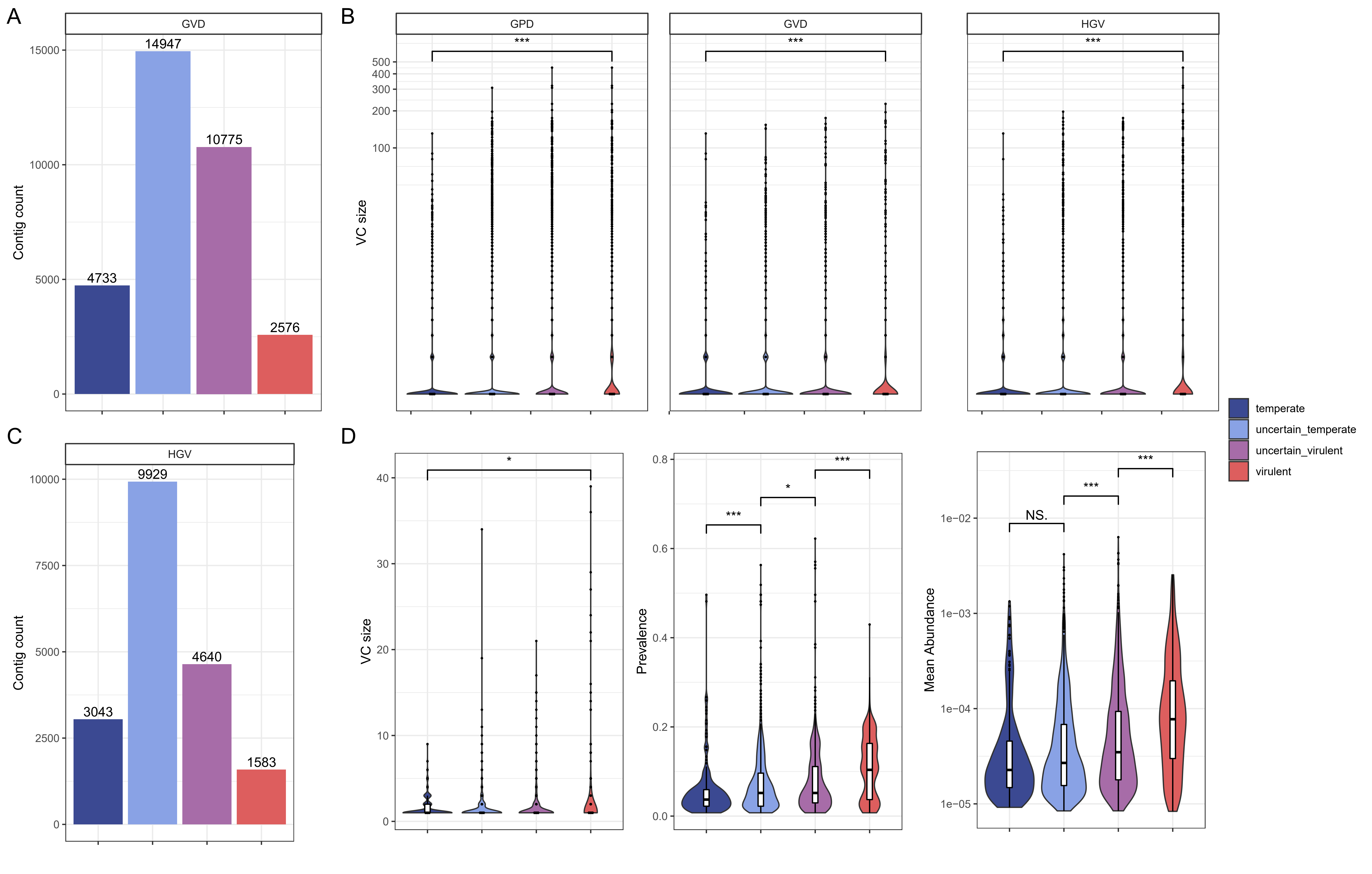
